# Benchmarking long-read genome sequence alignment tools for human genomics applications

**DOI:** 10.1101/2021.07.09.451840

**Authors:** Jonathan LoTempio, Emmanuèle Délot, Eric Vilain

## Abstract

**Background:** The utility of long-read genome sequencing platforms has been shown in many fields including whole genome assembly, metagenomics, and amplicon sequencing. Less clear is the applicability of long reads to reference-guided human genomics, the foundation of genomic medicine. Here, we benchmark available platform-agnostic alignment tools on datasets from nanopore and single-molecule real-time platforms to understand their suitability in producing a genome representation.

**Results:** For this study, we leveraged publicly-available data from sample NA12878 generated on Oxford Nanopore and sample NA24385 on Pacific Biosciences platforms. Each tool that was benchmarked, including GraphMap2, LRA, Minimap2, NGMLR, and Winnowmap2 produced the same alignment file each time. However, the different tools widely disagreed on which reads to leave unaligned, affecting the end genome coverage and the number of discoverable breakpoints. Minimap2 and winnowmap2 were computationally lightweight enough for use at scale. No alignment from one tool independently resolved all large structural variants (10,000-100,000 basepairs) present in the Database of Genome Variants (DGV) for sample NA12878 or the truthset for NA24385.

**Conclusions:** It should be best practice to use an analysis pipeline that generates alignments with both minimap2 and winnowmap2 as both are lightweight and yield different views of the genome. If computational resources and time are not a factor for a given case or experiment, a third representation from NGMLR will provide another view, and another chance to resolve a case. LRA, while fast, did not work on the nanopore data for our cluster, but PacBio results were promising in that those computations completed faster than Mininmap2. Graphmap2 is not an ideal tool for exploration of a whole human genome generated on a long-read sequencing platform.

## Introduction

The diverse ecosystem of DNA preparation, sequencing, and mapping technologies is capable of generating computational representations of genome biology through different chemistries and processes on different sequencing and mapping platforms [1–5]. This has resulted in many different approaches for making use of the data of different sources, with algorithms and the tools built upon them used across platforms to differing success. Here we briefly outline the differences between these platforms and analysis strategies so that we may consider an important gap in applying the newest technologies to the problem of human genomics.

Sequencing-by-synthesis platforms produce highly accurate sequence data in the form of short-reads (<300 basepairs, bp) [6] from high molecular weight DNA inputs. These molecular input libraries can be prepared for short read sequencing (SRS) that are oriented in context long-range genome position through high-throughput chromatin conformation capture (Hi-C) methods [7]. In contrast, single-molecule real-time platforms can produce highly accurate reads through circular consensus sequencing (CCS) on molecules >10 kilobasepairs (kbp) [8] while nanopore-based sequencing [9] and nanochannel-based mapping platforms [10] can sequence or visualize megabase-length DNA molecules [11]. Each of these platforms has different abilities and available tools, which contributes to the diversity of projects enabled by these technologies [12,13].

Optical genome mapping (developed by Bionano Genomics) excels at detecting structural variants (SV), such as balanced translocations and deletions/insertions in the 1 kb to 1 Mb range, and its clinical utility was demonstrated in Duchenne (DMD) or facioscapulohumeral (FSHD) muscular dystrophies [14,15], and cancer. But long-read sequence (LRS), developed by Pacific Biosciences and Oxford Nanopore Technologies among others, has been shown to be the most appropriate technology to detect smaller variants in the 50 bp-1 kb range [16]. This is especially true in repetitive regions of the genome and should therefore bring new diagnosis potential for diseases of trinucleotide repeat expansion and genome instability such as Huntington’s Disease and Myotonic Dystrophies [17–21], as well as for discovery of SVs affecting regulatory regions of known genes. Fulfilling the promise of LRS for medical genetics requires understanding the tools that power the technology and their respective strengths.

LRS platforms have been further shown to be capable of generating single, phaseable reads spanning repetitive or complex genomic regions that remain unresolved in all [8,9,22,23] but one current human genome assembly [24]. This has implications for resolution of highly homologous pseudogenes and large SV, even in low-complexity genomic regions, or from otherwise healthy diploid samples. Read lengths in the tens of kbp have further allowed for end-to-end viral genome sequencing [25], while read lengths in the millions of bp have the potential to span whole mammalian chromosomes [9,26]. This potential for more contiguous de novo human assembly has led to studies to specifically improve and benchmark basecalling and polishing tools [27], as well as assembly tools [28–30] that can be applied to human genomics.

For reference genome-guided experiments, LRS has proven useful for amplicon sequencing in cancer detection [31] and for metagenomics [32–34], but the field is still in early days for assessment of variation in high-coverage human whole genome sequence. Indeed, high-coverage human whole genome sequence on LRS platforms has yet to be undertaken on the scale of projects based on Illumina-developed SRS platform [35,36]. This difference in data availability is reflected in the number of tools for SV calling - including at least 21 for SRS and 9 for LRS [13,37–40]. It is certain that, due to the richness of the sequence data and diminishing cost of generating those data, these LRS platforms will be used in large-scale reference-guided whole human genome projects that are presently dominated by SRS data production [41].

The value of LRS in resolving SV has been demonstrated by efforts aggregating callsets across platforms and technologies to deeply characterize a genome [16]. These experiments are concerned mainly with understanding the full scope of the architecture of a genome, rather than contrasting differences between results of one alignment tool or another. Each of these tools showcase their utility and strengths in their initial publications, especially in terms of the mapping quality of reads to the reference genome and precision and sensitivity to preserve known variants in synthetic or downsampled genomic data. However, there have been no studies that specifically benchmark LRS platforms and tools for reference-guided experiments.

To address this gap, we benchmarked the most recent LRS alignment tools with the datasets generated from the Joint Initiative for Metrology in Biology’s Genome in a Bottle Initiative (GIAB), specifically samples, NA12878 sequenced with nanopore technology and NA24385 sequenced with CCS technology. These were generated with Oxford 9.4 pore chemistry and with Guppy5 basecalling performed by the Whole Genome Sequencing Consortium [9] and with Pacific Biosciences RSII SMRT CCS technology by PacBio and the National Institute of Standards and Technology [42]. We compared computational performance (peak memory utilization, central processing unit (CPU) time, file size/storage requirements), genome depth and basepair coverage, and quantified the reads left unaligned in any given experiment. We have limited this study to tools that are platform agnostic and function on our cluster. Since the resolution of large SVs is a key application of this technology, and also allows comparison of the differences in genome alignments in an aggregate way, we ran the SV-calling tool Sniffles to highlight differences in breakpoint location in each binary alignment map (BAM) file [43]. Taken together, these experiments present a comprehensive view of differences in the products of LRS whole-genome alignment pipelines.

## Materials & Methods

### 1. Tool selection criteria

a. Exclusion criteria were designed after a comprehensive literature review of alignment tools. Starting with tools that were recommended by the developers of each platform, we examined the tools that were cited or benchmarked against new, platform-specific tools. We also used the website longreadtools.org, searching their database with the filters “nanopore”, “pacbio”, and “alignment”. Since this yielded many software tools, we dove deeper and excluded any tools not designed for whole genome experiments. Tools which passed this test were then assessed for their suitability for use with nanopore and SMRT data and whether they produced a SAM/BAM file for analysis. Since tools can be regularly updated or out-versioned, we wanted to use only the most up to date software at the time of analysis.
b. In our examination of structural variant calling pipelines, we elected to limit our study to the variant caller sniffles due to the fact that the output files were most descriptive.

### 2. Data

#### Main experiments

Reference genome: GRCh38 was accessed on March 22, 2022 from ftp://ftp.ncbi.nlm.nih.gov/genomes/all/GCA/000/001/405/GCA_000001405.15_GRCh38/seqs_for_alignment_pipelines.ucsc_ids/GCA_000001405.15_GRCh38_no_alt_analysis_set.fna.gz

#### Sequence data

NA12878 data from the WGS Consortium, rel7 guppy 5 basecalls was accessed on March 22, 2022 from: https://github.com/nanopore-wgs-consortium/NA12878/blob/master/Genome.md15kb insert size SMRT CCS data:

NA24385 was accessed from the human pangenomics consortium github on March 22, 2022 from: https://github.com/human-pangenomics/HG002_Data_Freeze_v1.0 SV truthsets

Structural variant truthsets for NA12878, updated February 25, 2020, from DGV were accessed on March 31, 2022 from: http://dgv.tcag.ca/dgv/app/downloads?ref=GRCh37/hg19 [71]

Structural variant truthsets from Pacific Biosciences and the GIAB consortium were accessed on May 20, 2022 from: https://github.com/PacificBiosciences/sv-benchmark [70]

There is a limitation in that these variants were called with alignments to GRCh37. However, since this study is conserved with SV size, rather than location, within 1,000 and 10,000 bp windows we are confident that this is a minor limitation.

#### Pilot experiments (S4)

##### Reference genome

GRCh38.p12 was accessed and downloaded on April 8, 2019 from https://www.ncbi.nlm.nih.gov/assembly/GCF_000001405.38/ [81].

##### Sequence data

Nanopore sequence data were accessed on April 8, 2019, version rel5-guppy-0.3.0-chunk10k.fastq, from AWS Open Data as made available by the Whole Genome Consortium. SMRT sequence data were accessed on May 28, 2019, version sorted_final_merged.bam from the NCBI FTP.

https://github.com/nanopore-wgs-consortium/NA12878/blob/master/Genome.md [82]

ftp://ftp.ncbi.nlm.nih.gov/giab/ftp/data/NA12878/NA12878_PacBio_MtSinai/ [83]

##### Database of genome variants

DGV data were accessed on January 30, 2020 from http://dgv.tcag.ca/dgv/docs/GRCh38_hg38_variants_2016-05-15.txt [84]. The full set of variants was reduced to 11,042 variants confirmed to be present in NA12878. 14 of these variants were excluded as they were called on contigs present out of the main reference assembly contigs.

### 3. Hardware configuration

Computations were performed on the George Washington University High Performance Compute Center’s Pegasus Cluster on SLURM-managed default queue compute nodes with the following configuration: Dell PowerEdge R740 server with Dual 20-Core 3.70GHz Intel Xeon Gold 6148 processors, 192GB of 2666MHz DDR4 ECC Register DRAM, 800 GB SSD onboard storage (used for boot and local scratch space), and Mellanox EDR InfiniBand controller to 100GB fabric.

### 4. Whole genome alignment tool benchmarking and analysis

The following tool versions were used in the work presented in the main tables and figures.

1. v1.3.3. https://github.com/ChaissonLab/LRA\ [48]
2. v0.6.3 https://github.com/lbcb-sci/graphmap2 [47]
3. v2.24 https://github.com/lh3/minimap2 [49]
4. v0.2.7: https://github.com/philres/ngmlr [43]
5. v2.0.3 https://github.com/marbl/Winnowmap [50]

There versions of alignment tools used in the preliminary experiments of this project available in S4 included:

1. v0.5.2 https://github.com/isovic/GraphMapLink [55]
2. v2.16 https://github.com/lh3/minimap2 [49]
3. v0.2.7 https://github.com/philres/ngmlr [43]

All tools were run with default parameters and were flagged for nanopore or SMRT data based on the requirements of that experiment.

Computational metrics were printed from SLURM job records with seff and sacct with flags –format JobID, JobName, Elapsed, NCPUs, TotalCPU, CPUTime, ReqMem, MaxRSS, MaxDiskRead, MaxDiskWrite, State, ExitCode. Samtools 1.15.1 was used for all alignment manipulations, alignment read depth coverage calculations, and to extract unmapped reads (69). Samtools view-f 4 was used to generate bamstats files and python venn was used to compare the readnames across unmapped read files to assess the degree of overlap of unmapped reads across these subsetted alignment files. R 3.5.2 version Eggshell Igloo with the tidyverse packages were used for preliminary experiments on depreciated data shown in S4. Genome coverage was calculated with samtools coverage. Binary conversion values were used for bytes (1,073,741,824) and kilobytes (1,048,576) to gigabytes.

### 4. Data reshaping and visualization

As data were integrated across multiple sources and formats, they needed to be reshaped for comparison and visualization. For example, the multiple separators found in VCFs (commas and semicolons) are not directly usable in python pandas. Relevant dataframes from genome alignment files were reshaped with shell scripts and python scripts whose methodology and key intermediate files are available on our github [85].

### 5. Comparison of breakpoints

Structural variants (SV) were called with sniffles/1.0.11 with default parameters [43]. Only variants called on the main GRCh38 assembly were included. Python was used to bin and graph structural variants by SV Type. Intermediary files and scripts are available upon request. Data from sniffles VCFs and from DGV were subsetted by comparable fields including SV Type, SV Length, and Chromosome since a common format was unavailable for direct comparison. Figures were made with seaborn and matplotlib [86,87] as well as Microsoft Excel.

## Results

Rigorously annotated variants were drawn from the Database of Genomic Variants (DGV) [44] for NA12878 and from a curated public repository for NA24385 [45]. We present our results in four sections: 1) the tools that were included, 2) computational performance and benchmarking, 3) an analysis of aligned and unaligned reads, 4) an analysis of structural variation present in each alignment compared to a baseline.

### 1. Tools that passed the inclusion/exclusion criteria

Following a literature review of available alignment tools and search of long-read-tools.org [46], we established a set of inclusion criteria (see Material and Methods), which accounted for both types of LRS data (a tool must be able to handle both nanopore and SMRT reads), as well as the state of the field in terms of software updates (must not have been superseded by another tool). All relevant tools and their reason for exclusion are outlined in Table 1, with all tools annotated as platform agnostic and relevant to genome alignment included in S1.

**Table 1.**
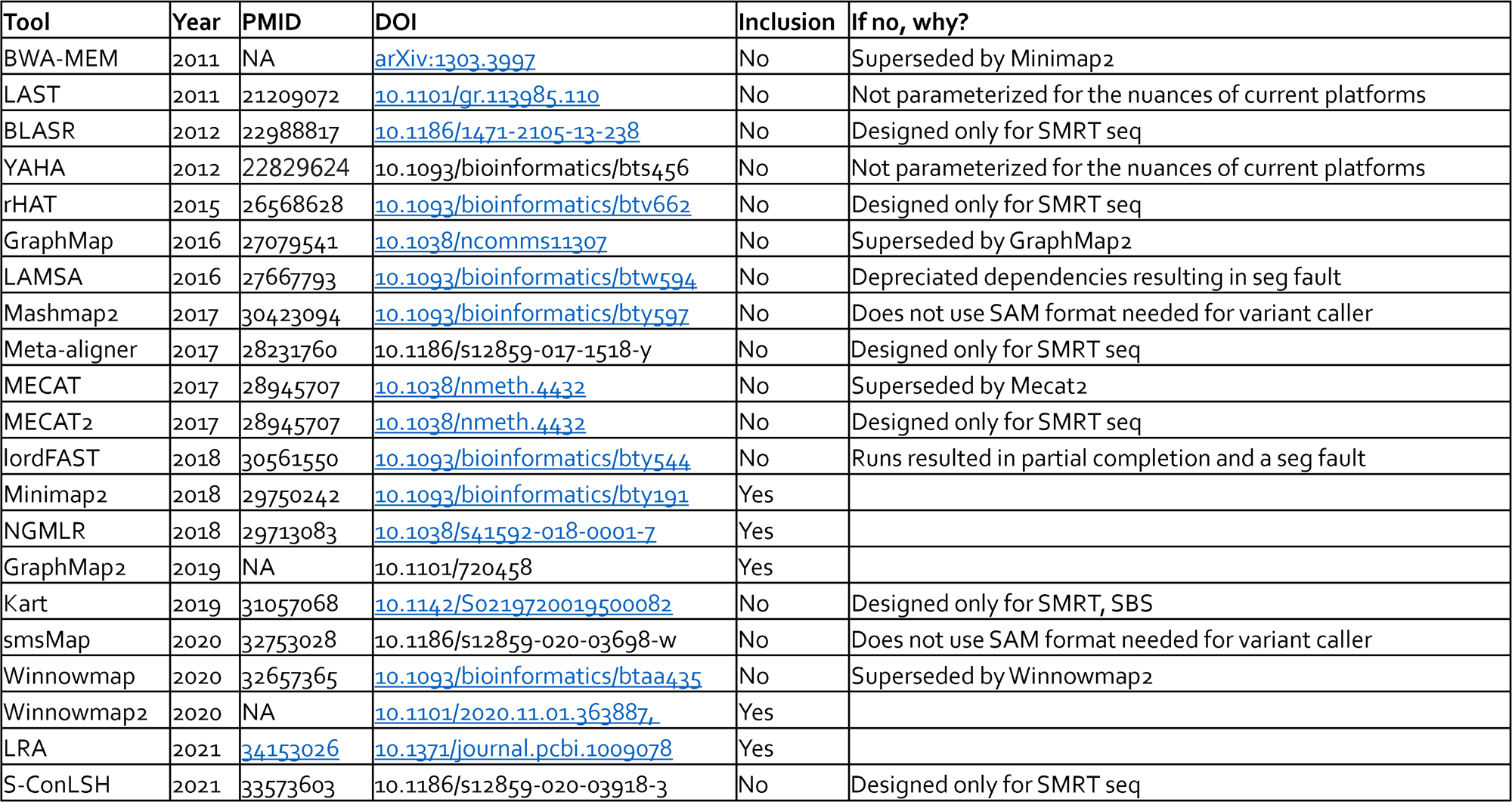
Long-read genome sequence alignment tools. The tools included in the study are highlighted in this study. Further tools from LongReadTools.org are available in S1.

Five alignment tools, GraphMap2 [47], LRA [48], minimap2 [49], NGMLR [43], and Winnowmap2 [50] were included in this study. Fifteen tools were excluded because they were superseded by other tools, produced an alignment file in a non SAM format, were designed for only one type of input data, or did not work on our cluster [51–64].

### 2. Computational performance and benchmarking

Computations were run three times on a single node allocated fully on our university’s high-performance compute cluster (HPC), which includes 40 CPUs with a configuration described in the methods. All tools were run with default parameters to reflect typical use in exploratory studies. High level computational benchmarks can be found in Table 2. Full reports on each run can be examined in Supplemental S2, while a composite table of the output of samtools coverage is available as Supplemental S3.

**Table 2.** Benchmarking metrics. Each of the three runs for each tool is shown in this table. All values are taken directly from slurm’s sacct and seff features. All metrics are intuitive except for CPU efficiency, which is a measure of idle CPUs to CPU time over wall clock time.

| Platform | Tool | Wall Clock Time<br>(D-H:M:S) | CPU Time<br>(D-H:M:S) | CPU<br>Efficiency | Memory Utilized<br>(Gb) |
| --- | --- | --- | --- | --- | --- |
| Nanopore | Graphmap2 | 14-00:00:04 | 560-00:02:40 | 65.25% | 90.19 |
|  |  | 14-00:00:17 | 560-00:11:20 | 63.34% | 94.58 |
|  |  | 14-00:00:15 | 560-00:10:00 | 64.23% | 89.68 |
|  | Minimap2 | 1-11:00:08 | 58-08:05:20 | 7.51% | 12.43 |
|  |  | 1-10:48:18 | 58-00:12:00 | 7.51% | 12.80 |
|  |  | 1-11:07:00 | 58-12:40:00 | 7.51% | 12.78 |
|  | NGMLR | 10-00:00:03 | 400-00:02:00 | 97.70% | 35.86 |
|  |  | 10-00:00:10 | 400-00:06:40 | 99.56% | 34.38 |
|  |  | 9-14:55:56 | 384-21:17:20 | 93.43% | 36.34 |
|  | Winnowmap2 | 15:25:58 | 25-17:18:40 | 83.82% | 29.48 |
|  |  | 15:19:14 | 25-12:49:20 | 83.61% | 30.02 |
|  |  | 15:15:24 | 25-10:16:00 | 83.06% | 28.74 |
| SMRT-CCS | Graphmap2 | 5-01:33:22 | 202-14:14:40 | 98.88% | 45.07 |
|  |  | 5-02:54:53 | 204-20:35:20 | 98.81% | 44.90 |
|  |  | 5-06:49:25 | 211-08:56:40 | 98.52% | 48.96 |
|  | LRA | 6:53:00 | 11-11:20:00 | 33.20% | 20.63 |
|  |  | 6:53:04 | 11-11:22:40 | 33.11% | 21.10 |
|  |  | 6:32:33 | 10-21:42:00 | 33.22% | 27.91 |
|  | Minimap2 | 11:02:04 | 18-09:22:40 | 7.51% | 9.10 |
|  |  | 11:06:03 | 18-12:02:00 | 7.51% | 9.13 |
|  |  | 10:55:09 | 18-04:46:00 | 7.52% | 9.09 |
|  | NGMLR | 21:50:29 | 36-09:39:20 | 99.20% | 25.53 |
|  |  | 22:24:38 | 37-08:25:20 | 99.19% | 25.69 |
|  |  | 21:58:53 | 36-15:15:20 | 99.53% | 25.82 |
|  | Winnowmap2 | 3:39:42 | 6-02:28:00 | 88.10% | 18.57 |
|  |  | 3:43:42 | 6-05:08:00 | 88.32% | 18.71 |
|  |  | 3:40:32 | 6-03:01:20 | 87.91% | 19.14 |

#### Globally successful tools

##### Minimap2 was the least memory-demanding tool

Minimap2 successfully aligned nanopore data every time with unrestricted and restricted resources. Unrestricted runs used around 12.6 gigabytes (Gb) and the jobs took ∼34 wall clock hours to complete.

Minimap2 also successfully aligned SMRT data every time with unrestricted or restricted resources. The runs used around 9 Gb and the jobs took ∼11 wall clock hours to complete, considerably faster than the (admittedly larger) nanopore dataset.

Memory usage and runtime were consistent across triplicate runs with unrestricted resources and did not change with restriction of resources when the tool was used with either dataset. The consistency of results, as well as the speed and relatively low computational demands of minimap2 make it a strong candidate for inclusion in clinical analysis pipelines.

##### Winnowmap2 was the fastest tool

Winnowmap2 was the fastest to run on any dataset. Specifically, it aligned the entire nanopore dataset in around 15 wall clock hours, using only a little more than two times more memory than minimap2. For SMRT, it was even faster at just over 3.5 wall clock hours and almost exactly twice as much memory. These numbers were consistent over all runs.

Additionally, the high CPU efficiency of the jobs on our cluster were positive to note: over 80% efficiency, relative to minimap2’s ∼7% efficiency. This is a measure of the ratio of CPU time used to the wall clock time times the number of CPUs. In the case of our cluster, CPUs are fully allocated to a node and not shared once allocated, but it does point the way towards either choosing more efficient tools, or optimizing jobs.

##### NGMLR completed 4/6 tasks and performance demanded great resources

NGMLR successfully aligned SMRT data every time but was the second slowest. The runs used around 25 Gb and the jobs took around 22 wall clock hours to complete, nearly twice as long as the next tool, minimap2.

NGMLR was much more inconsistent on nanopore data. It successfully aligned the data on our first range finding experiment in approximately ∼230 wall clock hours. Following this run, two jobs were set with a time of 10 days, and both resulted in timeouts. The single successful run used 36.34 Gb of memory, the most of all tools which produced an alignment.

#### Tools that presented a challenge

##### LRA did not work on nanopore-generated data

While LRA did show promise on SMRT data, aligning a genome in ∼11 CPU days or ∼6 wall clock hours, we could not run it successfully with nanopore. Each run resulted in unresolvable segmentation faults. In the case of SMRT data, it did fall in the front of the pack, aligning genomes quickly with typical memory usage relative to its peers (∼20-27GB)

##### GraphMap2 was the most resource-intensive tool

GraphMap2 took the longest, used the most memory, and had the most failures of all of the tools we examined. It did not produce alignment files on nanopore data in the time limit of 14 days imposed by our HPC. On the SMRT dataset which it was successful at aligning, it took more than 5 times longer to produce an alignment file than the next slowest tool, NGMLR. It had the highest memory usage, in excess of 89 GB nanopore on 44GB on SMRT.

### 3. Whole genome alignment

#### Aligned reads show disagreement in coverage of each chromosome

In preliminary experiments that completed successfully, alignments of the same genome with the same tool had the same genome coverage, whether resource-restricted or not (S4, column O). For this reason, one file generated from the first run was used for subsequent analysis, shown in Table 3, a summary of the coverage depth of each chromosome alignment and the percentage of the basepairs of GRCh38 covered. Global genome coverage of ∼30x was reported for nanopore [9] and ∼34x, 20x of which is present in the 15kbp insert set, was reported for SMRT [42,65,66] data sets. Breaking down the summary into chromosomes highlights allows for more precision than referring to a genome by its coverage as a uniform metric, since there is discrepancy across the chromosomes.

**Table 3.**
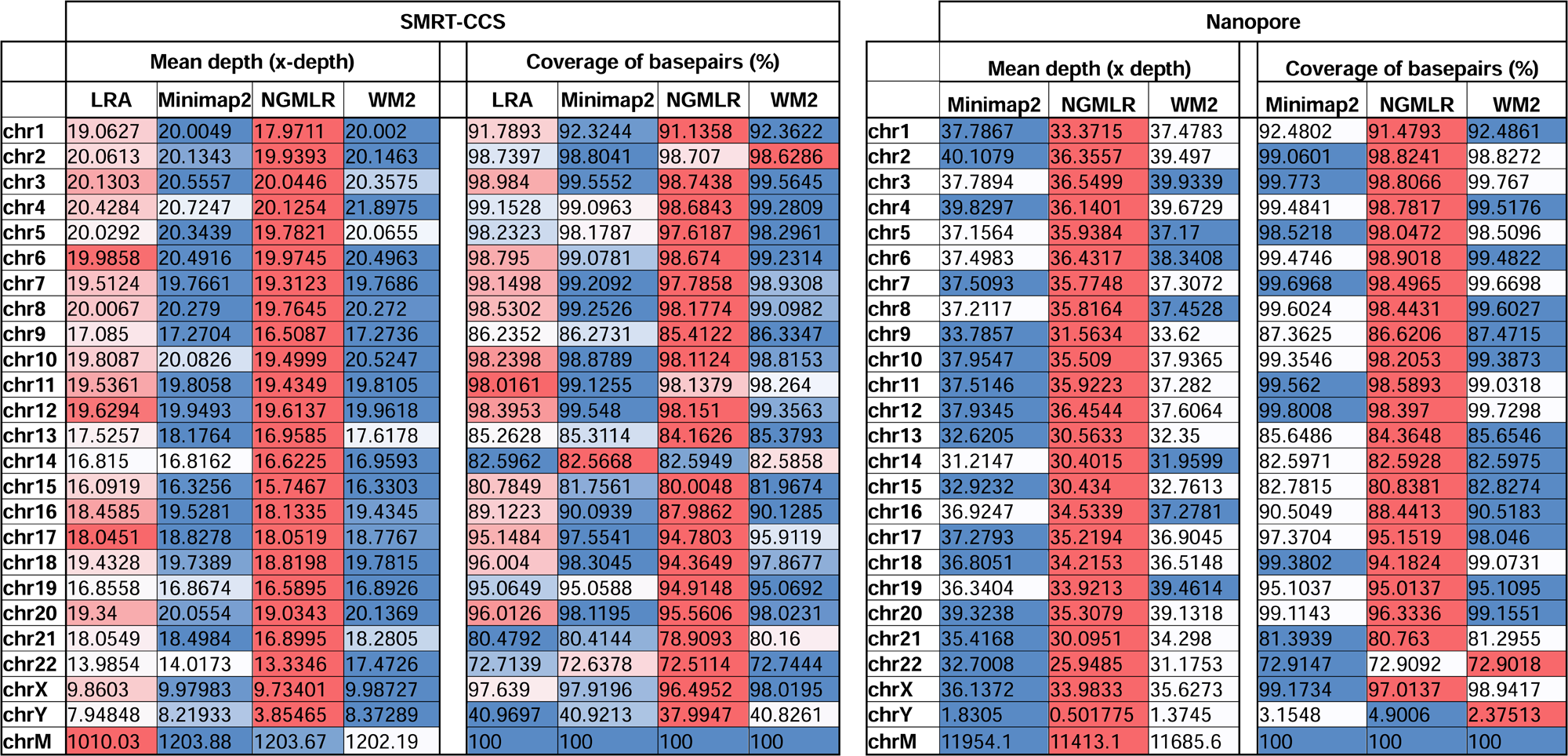
Chromosome level descriptions of the average number of reads covering each basepair, as well as the percentage of basepairs of the reference covered. Cells are hued within a row by chromosome to compare coverage on each chromosome in each dataset. Winnowmap2 abbreviated as WM2.

For nanopore data, three alignment filles were measured, while for SMRT data, four files could be included. Table 3 is hued horizontally by chromosome to highlight differences. It becomes immediately apparent that minimap2 and winnowmap2 both retain the most reads (highest x coverage) and cover the most basepairs of the reference. In plain language, they are the bluest columns. The column which represents NGMLR-aligned data is the reddest in all cases, suggesting that NGMLR is excluding the most reads and covering the fewest basepairs of the reference genome, which may impact downstream analysis. The final tool, LRA, which only worked on SMRT data, is intermediate with some light reds and whites in terms of coverage depth, and some blues in terms of coverage of basepairs.

#### Unaligned reads reveal differential exclusion of reads

The readname assigned to each read in a fastq retained in the BAM allowed us to directly compare the lists of reads that were not included in the alignment by each alignment tool with python venn (Fig. 1) [67].

**Figure 1.**
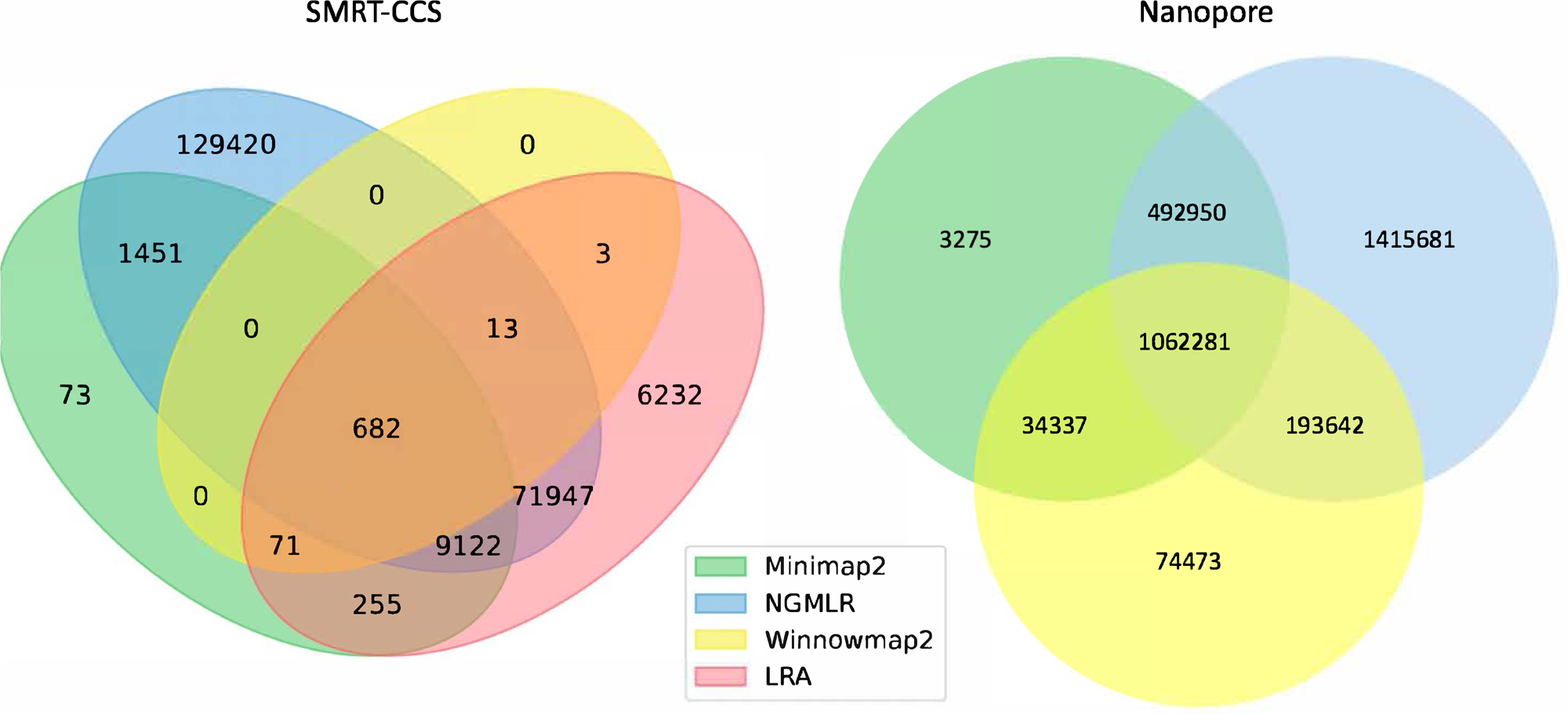
Unmapped reads. Venn diagrams show the total number of reads flagged as unmapped in the sorted BAM files. Panel A presents unmapped reads from the nanopore dataset. Panel B presents unmapped reads from a SMRT dataset.

All tools agreed to leave ∼1 million of the same nanopore reads unaligned, but differed in their overall totals of unaligned reads. NGMLR left the highest number of reads unaligned, ∼3.1 million, which explains why it produced the lowest coverage genome. It agreed with the other tools on approximately half of its discarded reads. Minimap2 and NGMLR agreed to leave a further ∼0.5 million reads unaligned, while NGMLR and Winnowmap2 agreed on a separate ∼0.2 million unaligned reads and Winnowmap2 and Minimap2 on ∼0.03 million further unaligned reads.

All tools agreed to leave 682 of the same SMRT reads unaligned. This is because winnowmap2 excludes less than 1000 reads in total, where the other tools excluded more reads over roughly three orders of magnitude. NGMLR left the highest number of reads unaligned, ∼200 thousand, which explains why it produced the lowest coverage genome. LRA excluded around 90 thousand reads in all, of which around 70 thousand are common to NGMLR. This also likely contributes to its lower-coverage status shown in Table 3. Minimap2 leaves out just shy of 12 thousand reads in all, most of which are shared with NGMLR and LRA.

In all, not enough information is available at this level of granularity to pass judgment on whether or not tools include or exclude the right reads. For this, an examination of breakpoints is required.

### 4. Breakpoints reveal differences between competing alignments of the same genome

By examining the coverage of the genome and the overlap of the discarded reads, it became clear that more detail was needed to understand the differences in alignment. To globally compare the alignments to each other, and assess their usability for variant calling, we looked to the SV known to be present in the NA12878 genome as curated by the DGV resource [44]. We ran a pilot study on now depreciated datasets for linear SMRT data [68] before undertaking a study with current technology. The results of these pilot experiments are available as S4 and point to a gap between the largest SV curated in DGV and what is resolvable by long-read sequencing.

To expand upon these initial findings, we accessed the most up-to-date data generated on nanopore and SMRT platforms. Presently, this includes the WGS consortium rel7 on NA12878 for nanopore and a new release of SMRT-CCS on NA24385. NA24385, the son of the Ashkenazi trio, is not present in DGV, so a high quality callset released by PacBio in collaboration with the Genome in a Bottle Consortium was used as a truthset [69].

We leveraged the sniffles SV caller because of its highly-detailed output files. Our truth sets were the curated SV present in DGV and the set released by PacBio and GIAB for NA24385. Due to the different nomenclature for SV type in the sniffles output and the NA24385 calls, which follow VCF specification [70] but differ from DGV annotations, comparisons were limited to the four classes of variant that were most unambiguously labeled: insertions, deletions, duplications, and inversions.

The variants available for NA24385 were coded as all insertions or deletions, with subtypes within including duplications. Therefore, the callsets from SMRT do not include inversions as there was no baseline. Sniffles variants were graphed by SV length on the X-axis in shades of blue, grey, and yellow, contrasted with DGV variants in red, organized by platform and SV type. Figures are organized by platform and size of SV, specifically between 1001 and 100,000 bp, the range at which LRS platforms are known to excel.

#### Nanopore

For the variants between 1,001 and 10,000 bp (Fig 2), alignment sets perform largely well on calling deletions that are curated in DGV while inversions were largely missed in all alignments. NGMLR alignments contained the highest number of breakpoints called as duplications. Above the 2001-3000 bp bin, the total duplications in DGV eclipses the number of breakpoints called in any of the alignment files. In contrast, there are very few insertions of this size range present in DGV, but hundreds in our callsets. This could be due to a differential interpretation of the breakpoints, where origin of inserted material has been ascertained (as a duplication) in DGV but remains unattributed (as an insertion) in call sets.

**Figure 2.**
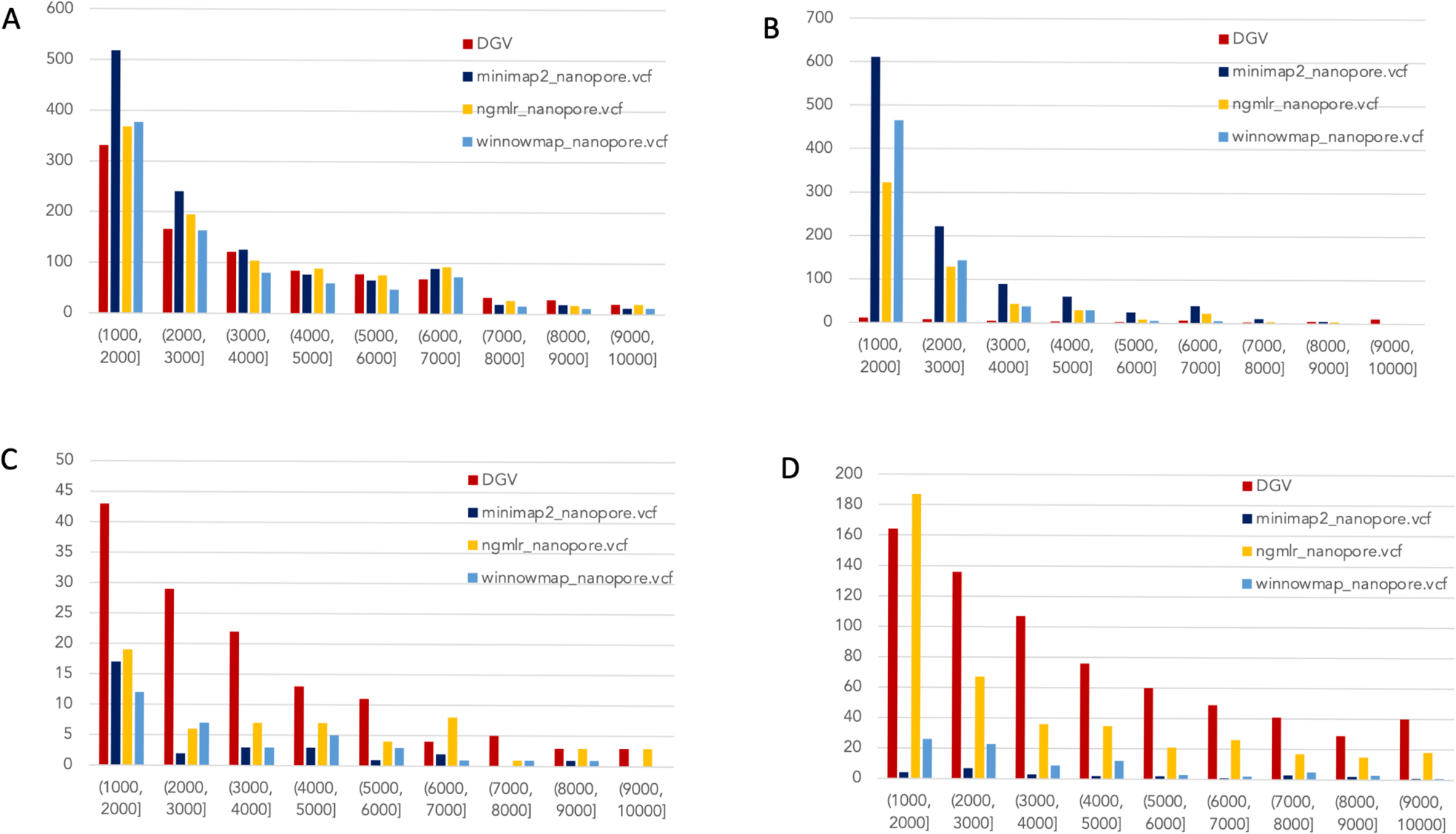
Nanopore SVs 1001-10,000bp. Variants called by sniffles on each alignment file. A. Deletions. B. Insertions. C. Inversions. D. Duplications.

In the 10,001-100,000 bp size range (S5), DGV largely contains more variants, with the exception of inversions. Inversions present an interesting case here because DGV contains more very large inversions than are reported (>80,001), but various callsets do well at smaller ranges. Across deletions, insertions, and duplications, all alignment tools fall short of the curated reference. It is most interesting that DGV had few insertions smaller than 10,000bp, but hundreds greater than that threshold. The trend of VCFs from our alignment files overperforming relative to DGV breaks down, and they highly underperform, calling only a few 10s of variants.

#### SMRT

The truthset for the SMRT dataset from NA24385 does not contain variants annotated as inversions, so Fig 3 (SV 1001-10,000bp) and S6 (SV 10,001-100,000bp) only contain three panels each. We can see immediately in Fig 3 that the bars generated from the reference set are much closer to the height of the bars from the VCFs, and this is not affected by the use of only the 15kbp insert set. There is one notable exception with regard to duplications. As with the nanopore dataset, an NGMLR alignment contains the most duplications.

**Figure 3.**
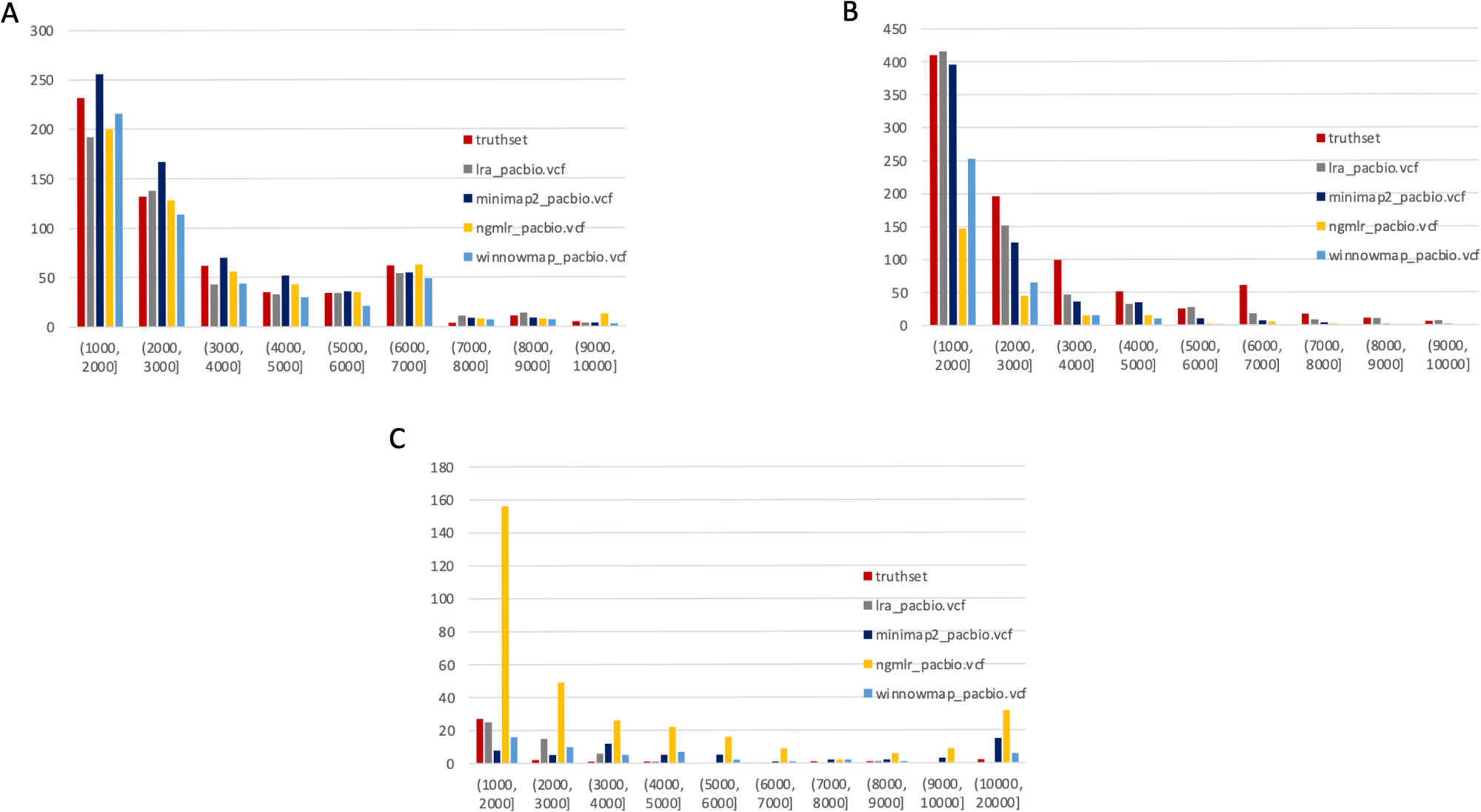
SMRT SVs 1001-10,000bp. Variants called by sniffles on each alignment file. A. Deletions. B. Insertions. C. Duplications.

However, these duplications are far in excess of what is present in the reference set. This is notable, as the reference set has been carefully validated. What was missed, versus what is a false positive, is not resolvable in this experiment and likely not resolvable without wet lab bench work or orthogonal validation.

The largest SV show a complex picture (S6). NGMLR calls the most and the largest deletions and no tools do well for large insertions in absolute terms or relative to the truthset. However, minimap2 and NGMLR preserve no breakpoints called as insertions. For duplications 10,001-20,000bp, all tools except LRA resolve many more than are present in the reference. NGMLR continues to include many more duplications than the other alignments, and more than the reference.

The disparity between the number of large variants greater 10,001bp in the NA24385 truthset versus the NA12878 is striking. This, along with the general lack of reference standards for all publicly-available genomes, highlights the need for better, more comprehensive reference sets for all publicly-available resources. Inability to resolve large duplications may yield false negative results for conditions where structural variation is a recognized etiology, such as DMD or FSHD, where variants range between the tens and hundreds of thousands of kbp [14,15], beyond the size of variants accurately detected in this study. It is also a critical issue when the technology is used for broad exploratory surveys seeking to identify new etiology in low-complexity regions of the genome where long-read sequencing should shine.

## Discussion

In this study we have highlighted the key differences between alignment files generated by different tools. By using well-characterized genome standards, NA12878 and NA24385, we were able to directly compare the performance of each tool on the sequencing datasets obtained on different platforms. We further analyzed the reads that were included or excluded from the alignment, and how these read alignments revealed breakpoints that could be resolved as structural variants in genome architecture. We then compared those variants to sets of previously published variants discovered through multiple platforms.

Reassuringly, each alignment tool was internally consistent: when an alignment tool was given the same fastq and the same reference genome, it produced the same result as judged by bamstats and sniffles variant callsets. However, when looking across the alignment files produced by different tools on the same sequence data, the representations of the genome diverged in terms of which reads were included or excluded and the numbers and types of variants that were present in VCFs. This is impactful because of the potential high value of LRS data in terms of genome phasing and identification of epigenetic DNA modification [71]. Since the majority of experiments that leverage large scale population surveys can be expected to rely on reference-guided alignment rather than de novo assembly because of both the cost and speed of analysis [72], it is key to understand the idiosyncrasies of each type of alignment files prior to generalizing clinical use for these technologies. Furthermore, clinical experiments in genomic medicine face human time constraints – speedier analyses will have higher appeal and adoption.

At this time, GraphMap2 does not show utility for producing whole genome alignments that include structural variations when run with default parameters. The resource usage was large and the time to complete the computation was long and when it did work, sniffles could not call variants on its product BAM. This is not unexpected, as the tool was designed in large part to increase single nucleotide variant sensitivity in noisy nanopore-sequenced reads [47,55]

Minimap2 used the least memory while winnowmap2 ran successfully in the fastest time, an important point, should these platforms be tied closer to bedside applications. On data from both platforms, they both allowed calling of the most insertions and deletions, but fell short on inversions and duplications.

NGMLR was the most discerning aligner, in that it left the highest number of reads unmapped. It used more compute power than Minimap2 and took much longer (3-10 times) than the next fastest tool. While it was designed specifically to resolve structural variation [43], it calls a high number of SV that have not been validated with other methodologies or curated in DGV.

There is a great divergence in sniffles-called variants from alignment files generated by all tools from the variants present in DGV. This is a concerning expansion of seminal findings in a previous study [16] as none of the sniffles VCFs mirror the SVs present in the high-quality curated DGV database. Furthermore, there are many genomes released across multiple consortia including GIAB and the Human Pangenome Reference Consortium [73]. But not all of these genomes are sequenced on multiple platforms, and comprehensive reference sets which are harmonious across samples, genomes, and datasets are still limited.

This resulted in us using two reference sets for this project to accommodate the latest releases of data. The callsets generated in our experiments used annotations which were different from the NA12878 variants in DGV and the curated variant set for NA24385, which was surprising. Finally, the most up to date variant benchmark for NA24385 is called on GRCh37-aligned genomes, which is from 2009 and not as comprehensive as current builds [24,74]. Taken together, this underscores the need for orthogonal approaches and collaboration between wet and dry labs to solve this problem.

The points above are critical in designing pipelines for genome analysis and structural variant discovery. In short, it is not a high burden to generate two alignment files per genome with each minimap2 and winnowmap2. In computationally unlimited research settings, there is value added in the generation of alignment files from NGMLR as well, although the time to complete these experiments makes them less appealing. These three perspectives on the same genome will account for some of the inherent differences of each tool and the algorithms they use to handle read alignment [75]. If computational resources are limited, minimap2 is the best choice to move the greatest number of genomes through the pipeline quickly with small memory needs; however, the loss of comprehensiveness must be considered in cases where a suspected variant is not found.

This is impactful in genomic medicine. For example, as variants range from tens of kbp in FSHD and hundreds of kbp to mbp in DMD, diagnosis of these disorders will likely not benefit from data generated on LRS platforms at present, underscoring the need for optical mapping or array-based technologies. However, disorders resulting from smaller SVs such as Huntington’s (∼18-540 bp) [76], myotonic dystrophy 1 (∼15-153 bp) and myotonic dystrophy 2 (∼338-143,000 bp) [77] (61) could be good candidates for deep study with LRS platforms based on the variants present in alignment files from minimap2, winnowmap2 and NGMLR. Accordingly, LRS has been used to identify variants in many such disorders [21,78].

If pathogenic loci are known, a high diagnostic yield may be obtained by generating maps with each available alignment tool, and use of a structural variant caller such as NanoSV [79]. Unlike sniffles, which provides a call of type of SV (deletion, insertion etc.), NanoSV only identifies breakpoints in the alignment, without assigning those breakpoints an SV type. A robust comparison of SV callers on nanopore datasets highlighted the relative strengths of variant calling pipelines and may help users determine the best caller for their experiments [80]. NanoSV may be suitable for identifying breakpoints missed by Sniffles, but comes with the further caveat that it is resource-intensive and may not scale in a clinical setting without vast computational resources [80].

Answers to biological questions can also leverage tools that work with data upstream and downstream of whole genome alignments; indeed, alternate solutions to discover differences in genome structure have been proposed that leverage new tools. Tools designed to specifically identify triplet repeat expansions in LRS data sets, such as Repeat Hidden Markov Model (Repeat HHM) or Tandem-Genotypes, have shown the superiority of the approach compared to other techniques [19,20]. RepeatHMM uses raw, unaligned reads in alternate ways than whole genome alignment to a reference [19]. Rather than aligning LRS, reads can be analyzed independently for microsatellites with specialized tools. The tandem-genotypes tool makes use of a LAST whole-genome sequence alignment to detect copy number changes in the genome [20]. In line with this fact, LRS has been used to identify variants in many disorders [21].

The discrepancies between the VCFs generated from alignment files starkly show the need to design experiments with the appreciation that the genomics ecosystem cannot yet be dominated by one platform or pipeline and requires a multifaceted approach to discovery. We are at a position where simply because the breakpoint is missing from the VCF from an LRS genome, we cannot say that it is not present. We must therefore look across platforms and data types for comprehensive genome representations [16]. However, this requires more data from the community - generation of NA24385 data on new nanopore chemistry would allow for direct comparison to the current CCS standard, or alternatively new CCS data could be generated on NA12878. This would address a critical gap in our ability to benchmark new tools in a platform-agnostic manner.

## Conclusions

As the cost of long-read sequencing catches up to that of inexpensive short read sequencing, the inevitable boom in data production will require well thought-out analysis pipelines. Pipeline design always involves a set of tradeoffs. To accurately assess these tradeoffs, we must have a rigorously benchmarked view into the tools available to create the analytic product. Here, we looked at the differences in reference-guided human genome alignments to understand the difference in each tool’s alignment of the same genome, and how it affects a structural variant callset. This informs our conclusion that, regardless of sequencing platform, when computational resources are not a limiting factor, it should be best practice to align an LRS human genome with minimap2, winnowmap2, and NGMLR to gain better insight into the architecture of a genome of interest. When compute resources are limited, minimap2 is a strong choice, and when time is a limiting factor, winnowmap2 is the best choice.

## Supporting information

Supplemental Table of Contents

## Acknowledgements

Adam Kai Leung Wong, PhD, High Performance Computing Specialist for Genomics at the GWU Computational Biology Institute provided critical support in preparing the HPC for this study.

