## Supplemental Table of Contents for "Benchmarking long-read genome sequence alignment tools for human genomics applications"

S1: Long-Read-Tools.org search results annotated for exclusion rationale

S2: Raw computational metrics from sacct and seff, including 3 tabs

S2_1: Summary of metrics

S2_2: Raw print from sacct

S2_3: Raw print from seff

S3: Data underlying coverage in Table 3, including 7 tabs

S4_1: PacBio Minimap2

S4_2: PacBio Winnowmap2

S4_3: PacBio NGMLR

S4_4: PacBio LRA

S4_5: Nanopore Minimap2

S4_6: Nanopore NGMLR

S4_7: Nanopore Winnowmap2

S4: Pilot benchmarks on depreciated datasets, including 3 tabs

S3_1: Computational metrics

S3_2: BAMstats

S3_3: Summary of sniffles-called and DGV-annotated SV

S5: Nanopore SVs 10,001-100,000bp.

S6: SMRT SVs 10,001-100,000bp.
