## Supplemental Files 5 and 6 for "Benchmarking long-read genome sequence alignment tools for human genomics applications"

### S5: Nanopore SVs 10,001 – 100,000 bp

#### Deletions

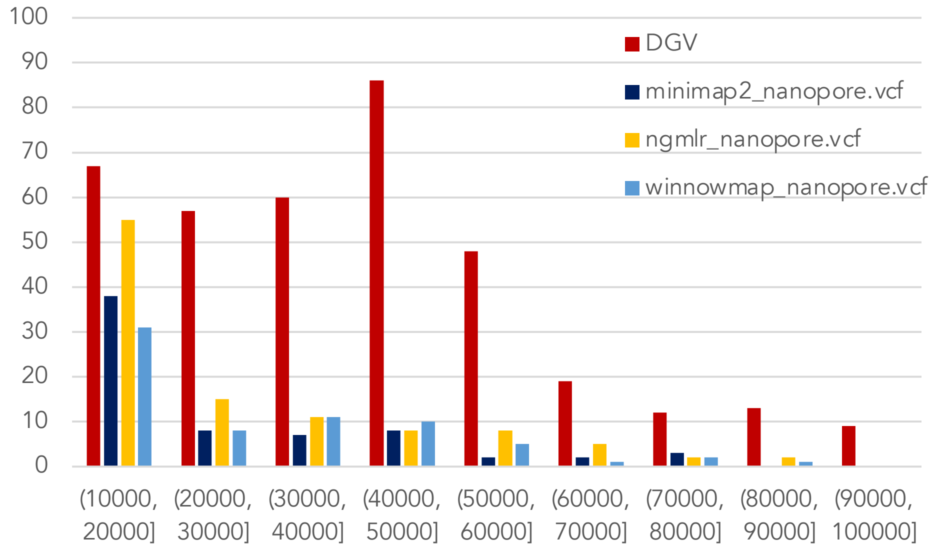


#### Insertions

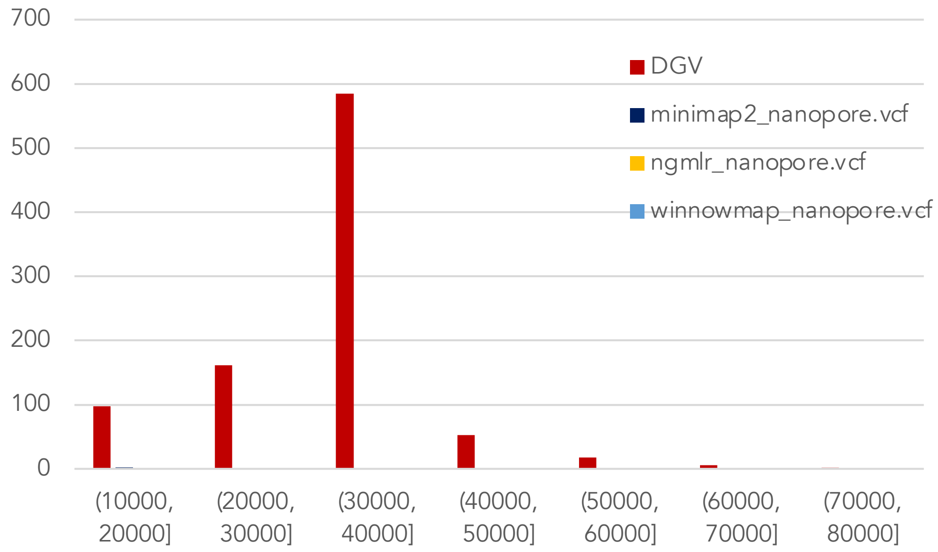


#### Inversions

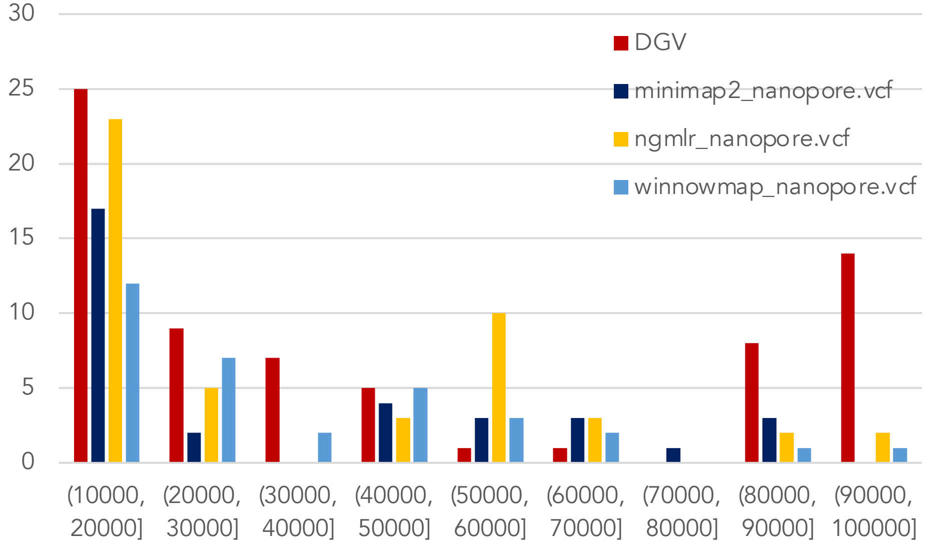


#### Duplications

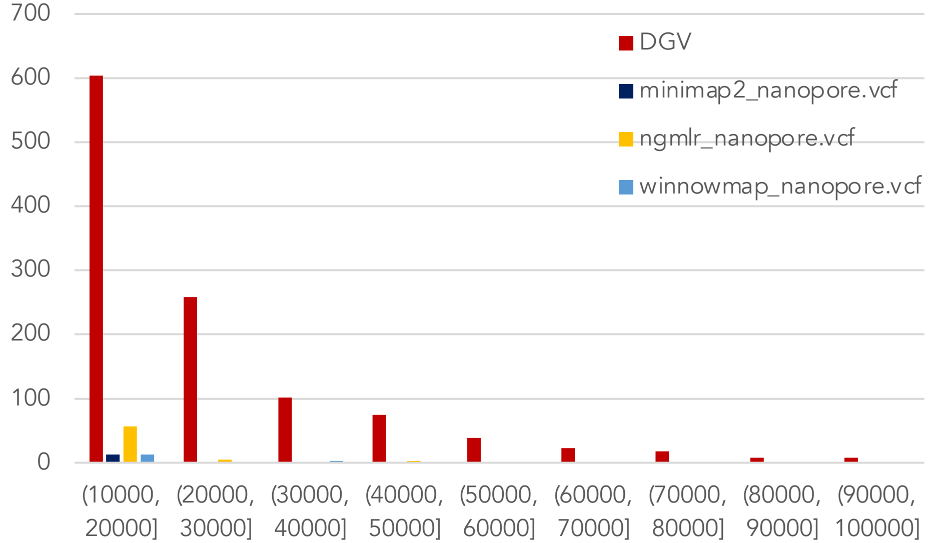

### S6: SMRT-CCS SVs 10,001 – 100,000 bp

#### Deletions

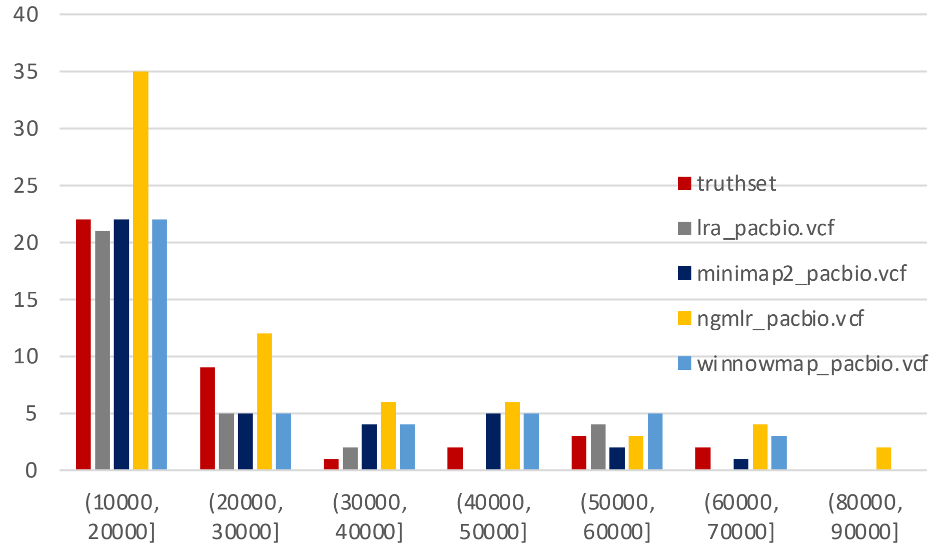


#### Insertions

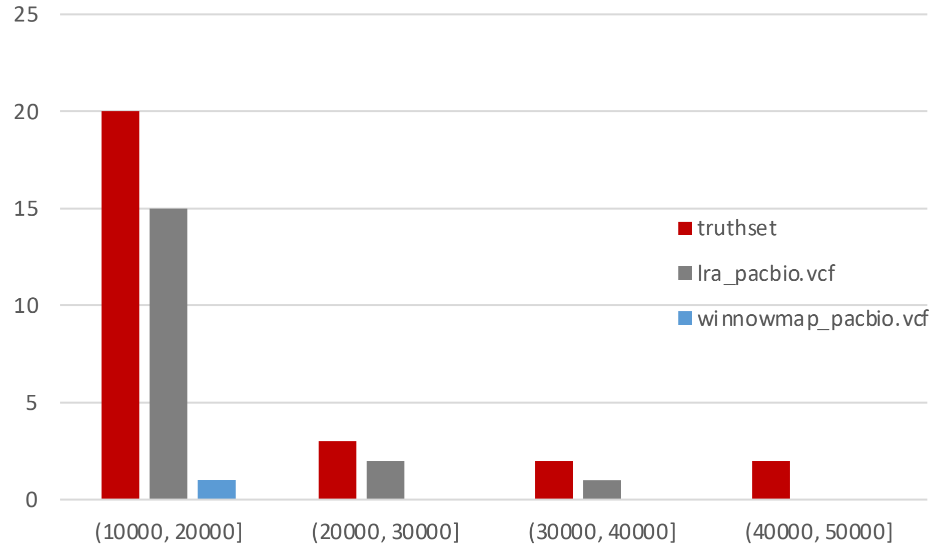


#### Duplications

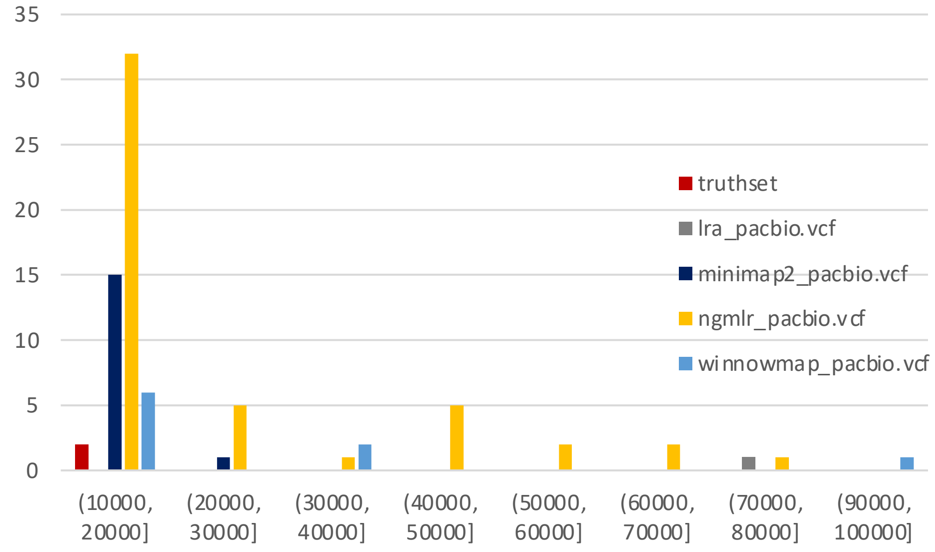
